# The integrity of perineuronal nets in the amygdala as a key factor in the resilience to social isolation stress in old mice

**DOI:** 10.1101/2023.08.04.551957

**Authors:** José Hidalgo-Cortés, Aroa Mañas-Ojeda, Francisco E. Olucha-Bordonau, Clara García-Mompó, Esther Castillo-Gómez

## Abstract

Major depression is the most prevalent neuropsychiatric disorder in elderly population, affecting more than 20% individuals over 60 years old, especially women. In this age range, social isolation is a major risk factor for depression. While there is a significant positive association between social isolation and depression in the elderly population, the neurobiological basis of this association is complex and still poorly understood. Evidence from animal models and human studies indicates that neuroplasticity, especially that of limbic brain regions, is impaired in depression but, till date, scarce studies address this question in older population. In this regard, animal models devoid of human cultural connotations represent a crucial tool. In the present study, we investigated the impact of chronic isolation stress (CIS) and a subsequent resocialization period in aged male and female mice (∼ 21 months-old), focusing our attention on affective symptoms and the plasticity of parvalbumin-expressing (PV+) neurons in the lateral/basolateral amygdala (LA/BLA). We found that CIS impaired affective behaviour and LA/BLA plasticity only in females. Specifically, CIS induced depressive-like symptoms and decreased the integrity of perineuronal nets (PNN). Resocialization effectively rescued all these impairments. Old males were not affected by CIS but in social conditions showed higher PNN integrity (less plasticity) than females. All together, our results demonstrate that old females are less resilient to CIS than old males and point to the integrity of PNN in the LA/BLA as a key regulator of depressive-like symptoms induced by social isolation.

## Introduction

Depression is the most prevalent mental disorder in the elderly population, even above dementia. It affects more than 20% of older people, especially women, that are 1.5-3 times more likely to suffer from depression than elderly men [1–3]. Several risk factors have been associated with depression, but stressful life events are thought to be especially relevant to the development of this disease [4–6]. In elder age, social isolation (or loneliness) is the most frequent stressor [7, 8], affecting between 30 to 50% of the senior population [9, 10]. Indeed, recent clinical studies have demonstrated a significant positive association between social isolation and depression in senescence [6, 11–14]. However, current treatment strategies for depression among elderly people have important limitations, in part because the neuropathological mechanisms of the disease in later-life are still poorly understood.

In the adult population, evidence from animal models and human studies indicates that the changes in neural plasticity induced by stress and other negative stimuli, especially in brain regions involved in emotional processing, play a significant role in the onset and development of depression [15, 16]. However, till date, scarce studies address this question in old population [17, 18]. Among all regions involved in depression, the lateral/basolateral amygdalar complex (LA/BLA) seems to play a pivotal role [19–21]. In fact, a variety of studies have been demonstrated morphological and functional changes of the LA/BLA associated with depression, including hyperactivity, hypertrophy and increased synaptic and structural plasticity ([22, 23] for review). Glucocorticoid receptors are highly expressed by LA/BLA neurons, including GABAergic neurons, what makes them one of the main targets of the neuroendocrine mediator of stress, the hypothalamic pituitary adrenocortical (HPA) axis [24, 25].

Around 50% of the GABAergic population in the LA/BLA are parvalbumin-expressing (PV+) fast-spiking interneurons [26]. These neurons reach a mature phenotype with the formation of perineuronal nets (PNNs) [27], which are specialized aggregates of extracellular matrix components (ECM) that enwrap their soma and proximal dendrites [28–30] and enhance their firing rate [31]. Accumulated evidence demonstrate that PNNs negatively control neuronal plasticity [32–34] by forming a physical barrier against new synapse formation while increasing the stability of the existing ones [35–38]. Perisomatic synapses are mainly stablished in the holes of the net-like structure of PNNs that are occupied by excitatory or inhibitory synaptic boutons [38, 39]. Notably, a recent study suggests that the integrity of PNNs is regulated by the size of their holes, with smaller holes stabilizing existing synapses and larger ones leaving “free” space for newly generated synapses, thus increasing plasticity [39]. PNNs are prone to degradation by ECM remodeling enzymes, specially the matrix metalloproteinase 9 (MMP-9), that actively participates in the disintegration of PNNs [34, 40]. Interestingly, the composition and expression patterns of PNNs in the brain, along with MMP-9 levels, are variable throughout life in a region- specific manner and according to level of plasticity of that region and that age [40–43]. In fact, PNN and MMP-9 interactions are so important for brain plasticity and function that abnormalities among them have been described in many neurological and psychiatric disorders including Alzheimer’s disease [44], schizophrenia [45] and major depression [46], among others [47, 48].

Available evidence suggest that PNNs in the amygdala are vulnerable structures to the effects of stress, which disrupts this “protective barrier” of PV+ interneurons, leading to increased instability within the circuitry [5]. Although there are many studies describing that such effects of stress on PNNs are dependent on age, and brain region [49], till date, there are no preclinical studies addressing this question in the LA/BLA of aged individuals. Likewise, no attention has been paid on the differential vulnerability of aged males and females to the effects of stress due to social isolation. Additionally, no study has addressed the question on whether a resocialization therapy can be effective in the recovery of the behavioral and neuronal alterations of aged mice. Therefore, in the present study we have investigated the impact of chronic isolation stress (CIS) and a subsequent resocialization period (CIS+R) in affective behavior and brain plasticity of aged male and female mice. We have hypothesized that CIS will be associated with depressive-like behavior and will impair the plasticity of PV+ neurons in the LA/BLA, by disrupting its protective barrier (PNNs). We have also analyzed whether these putative changes are dependent on sex and whether they can be rescued by resocialization. We believe that this and future studies on the mechanistic link between PNNs and depression in aged population may be key to develop novel and more effective drugs and therapies for loneliness-related depression in senescence.

## Materials and Methods

### Animals and experimental procedure

C57BL/6J mice were purchased from The Jackson Laboratory (Bar Harbor, Maine, EEUU) and bred in our animal facility (Animal Experimentation Service, Universitat Jaume I de Castelló, Spain). After weaning, male and female pups were separated by sex and housed in groups of 3-4 animals in conventional Makrolon® EU Type 2L cages (floor area: 530 cm^2^). All animals were maintained in a controlled environment with 12h:12h light-dark cycle (lights on from 8:00 AM to 8:00 PM), room temperature at 21 ± 1°C, humidity of 50 ± 5% and food and water available *ad libitum*. At old age (20.8±0.2 months old), mice were randomly assigned to each experimental group as described below (experimental procedure, **Fig. 1A**). Experiments were conducted in accordance with the guidelines provided by the European Community Council Directive (2010/63/EU) for the use of laboratory animals and were approved by the Committee on Bioethics of the Universitat Jaume I.

**Figure 1.**
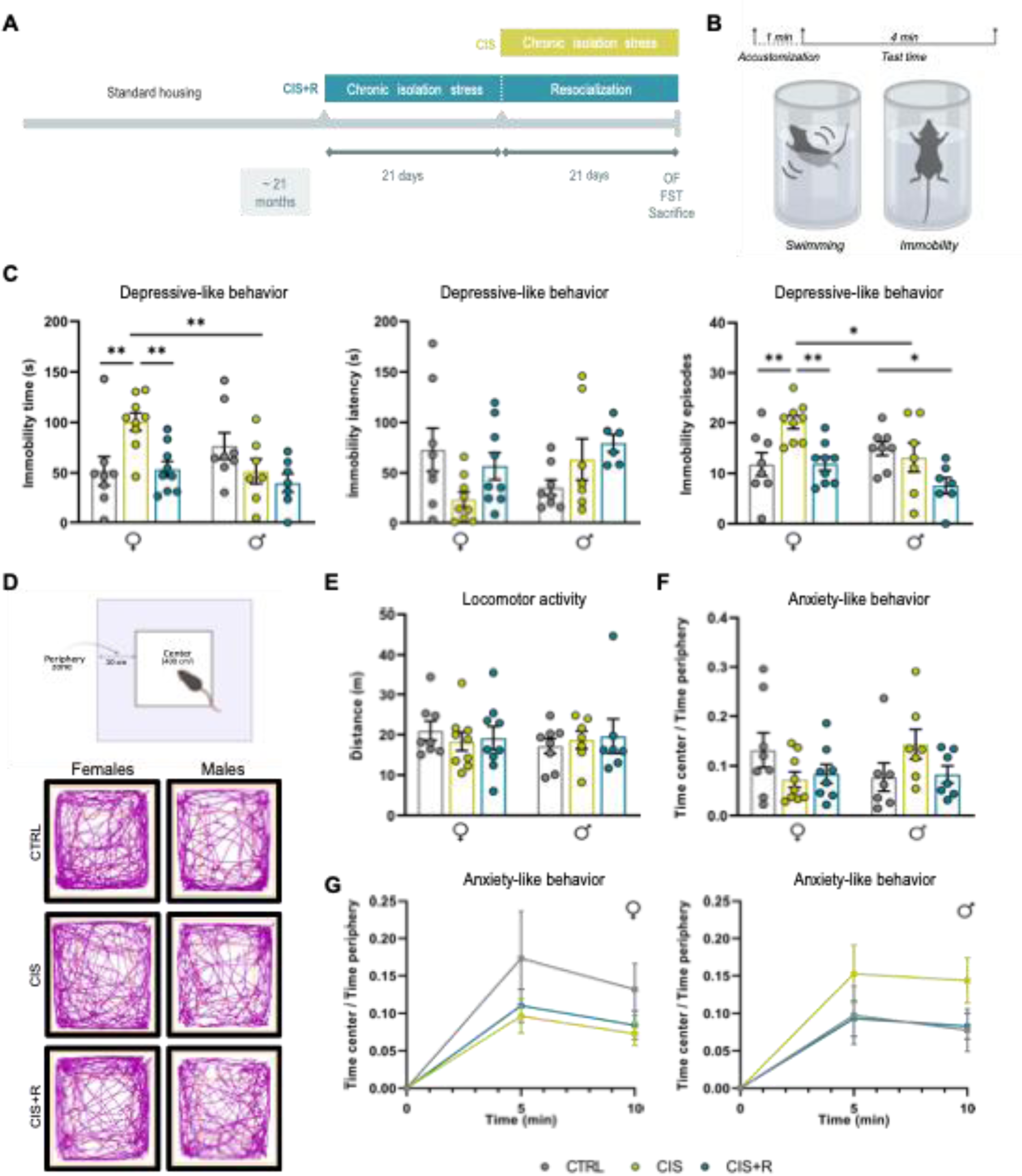
Affective behavior measured in the forced swimming test (FST) and open field test (OF). (A) Schematic representation of the experiment timeline. (B) Graphical explanation of the FST procedure. Upper panel: timeline of the test with the accustomization period (1 min) and test period (4 min). Lower panel: drawings indicating swimming state (left) and immobility state (right). (C) Immobility time (s), immobility latency (s) and immobility episodes were measured asan indicator of depressive-like behavior. (D) Schematic representation of the OF paradigm indicating the periphery zone and the center zone (upper panel) and representative track plots of females and males of each experimental condition (lower panel). (E) Locomotor activity was calculated by measuring total distance traveled in 10 minutes. (F) The ratio of time spent in the center to time spent in the periphery of the OF was calculated as an indicator of anxiety-like behavior. (G) The ratio Time center/ Time periphery between 0-5 min of test and 5-10 minutes of test is also plot. Data in graphs are presented as mean ± SEM (n=7-9/group) and asterisks represent statistically significant differences among groups, p<0.05 (*), p<0.01 (**). Each dot represents each animal. OF, Open Field; FST, Forced Swim Test; CTRL, Control group; CIS, Chronic isolation stress; CIS+R, Chronic isolation stress + resocialization.

### Social isolation and resocialization protocol

Old male and female mice were randomly divided into 3 groups for each sex: (1) chronic isolation stress (CIS), (2) chronic isolation stress + re-socialization (CIS+R) and (3) control group (CTRL) (**Fig. 1A**). No changes in housing conditions were applied to the CTRL group mice, so they remained socially housed in Type 2L cages with the same 3-4 animals with which they had been housed since weaning. Animals in the CIS group were individually housed for 21 days in conventional Makrolon® EU Type 2 cages (floor area: 375 cm^2^). CIS+R animals were socially isolated for 21 days as described for the CIS group and, afterwards, they were returned to their original home cage for 21 days (re-socialization with their previous cage mates). At the end of the experiment, all mice were tested for depression-like behavior (forced swim test, FST; **Fig. 1A-B**), locomotor activity and anxiety-like behavior (open field, OF; **Fig. 1A, D**). After behavioral testing, females underwent a vaginal smear (to check for persistent vaginal cornification associated to senescence) and then they were transcardially perfused. Both males and females were perfused approximately 45-60 minutes after the end of the behavioral tests (**Fig. 1A**).

### Vaginal smears

In order to confirm the persistent vaginal cornification associated to senescence [50, 51], vaginal smears from all females included in our study underwent microscopic examination. Each sample was obtained by flushing three to five times the vagina with saline solution (NaCl 0.9%, Sigma-Aldrich, St. Louis, MO). The final flush was placed on a glass slide, air-dried, and stained with toluidine blue. Then we calculated the proportion of leukocytes, nucleated epithelial cells and anucleated and keratinized cornified cells observed under the microscope, as described in [52–54]. All females in our study presented vaginal cornification, defined as the presence of >70% of cornified cells (data not shown) [50, 51].

### Behavioral tests

#### Forced Swim Test (FST)

Animals were individually tested inside clean glass beakers (30 cm tall x 15 cm Ø) half filled with water at 24±1°C. At the start of the test, each mouse was gently placed in the center of the beaker and was allowed to swim freely for 5 min (**Fig. 1B**). Animal behavior was video-tracked and analyzed by ANY-maze software (ANY-maze video tracking system v7.20; Stoelting Europe). Trials lasted 5 minutes, but behavior was only scored for the last 4 minutes (test time), except for the immobility latency (time spend until the first immobile episode). Immobility was defined as the absence of any movement, except of those that are necessary to keep the head above water. Immobility latency, time immobile and immobility episodes were considered as indicators of depressive-like behavior [55, 56]

#### Open Field (OF) Test

The OF apparatus consisted in a black opaque plexiglass covered arena (40 × 40 cm). At the start of the test, each mouse was placed in the center of the arena and was left to freely explore it for 10 min. The movement of the animal was video-tracked and automatically analyzed by ANY-maze software (ANY-maze video tracking system v7.20; Stoelting Europe). Ethanol 70% was used to wipe the chamber between tests. For analysis, the center and periphery zones were defined offline but were not marked on the apparatus to prevent the animal from stopping to explore the marks. The periphery zone was demarcated as the area located between 0 and 10 cm from the walls of the apparatus and the center zone was the remaining 400 cm^2^ area located at the middle region of the arena (**Fig. 1D**). The total distance travelled was considered as an indicator of locomotor activity and the time spent in the center zone versus the time spent in the periphery zone was considered as a parameter related to anxiety-like behavior as described before [57, 58]. For this last parameter, we also analyzed the potential differences between the first 5 min of the test (novelty phase) and the last 5 min of the test.

### Perfusion and microtomy

Prior to intracardial perfusion, animals received an overdose (120 mg/kg, i.p) of sodium pentobarbital (Dolethal, 200 mg/ml, Vetoquinol S.A., Madrid, Spain). After losing all reflexes, the animals were perfused first for 5 min with NaCl 0.9% and then for 15 min with 4% paraformaldehyde (PFA, Sigma-Aldrich, St. Louis, MO) in sodium phosphate buffer (PB 0.1 M, pH 7.4; Sigma-Aldrich, St. Louis, MO). Thirty minutes after perfusion, brains were removed from the skull and left 2h in PFA solution for post-fixation. Brains were then rinsed in PB 0.1M and cryoprotected with 30% sucrose in cold PB 0.1 M (4 °C) for 48 h. Cryoprotected brains were cut in 50-µm-thick coronal sections using a freezing sliding microtome (Leica SM2000R, Leica Microsystems, Heidelberg, Germany). Slices were collected sequentially in six subseries and stored until use at -20°C in Eppendorf tubes with cryoprotective solution (30% ethylene glycol, 30% glycerol and 40% PB 0.1M; Sigma-Aldrich, St. Louis, MO).

### Multiplexed immunofluorescence

Sections were processed free-floating for multiplexed immunofluorescence following the protocol we described previously [59]. Briefly, sections were washed in PBS and incubated for 1 hour with 10 % normal donkey serum (NDS; Biowest LLC, Kansas City, USA) in PBS with 0.2% triton-X100 (PBST, Sigma-Aldrich, St. Louis, MO). Then, sections were incubated with polyclonal rabbit IgG anti-PV (1:10000 Swant, Burgdorf, Switzerland), biotin-conjugated *Wisteria floribunda agglutinin* (WFA; 1:200 Sigma-Aldrich) in 5% NDS and PBST for 48h at 4°C. WFA is commonly used in brain tissue sections to detect perineuronal nets (PNNs) [60]. After being rinsed, sections were light-protected and incubated at room temperature for 2 h with donkey anti-rabbit IgG AlexaFluor®488-conjugated (1:400; Jackson ImmunoResearch, Pennsylvania, USA) and Streptavidin AlexaFluor®594-conjugated (1:200; Jackson ImmunoResearch, Pennsylvania, USA) in 5% NDS and PBST. All sections were mounted on gelatin-coated slides and coverslipped using fluorescence mounting medium (Fluoromount-G, ThermoFisher, Massachusetts, USA).

### Confocal microscopy and image processing

#### Densitometric analysis of PV+, PNN+ and PV+PNN+ neurons in the LA/BLA

Images covering the whole extension of the LA/BLA were obtained at 10x magnification in a Leica SP8 laser scanning confocal microscope (Leica Microsystems, Wetzlar, Germany). Z-stacks covering the whole depth of the sections were taken with 1 μm step size and only subsets of confocal planes with the optimal penetration level for PV and WFA were selected (**Fig. 2**). Counting was performed manually in all selected Z-planes using FIJI software (ImageJ, Maryland, USA) by a researcher blinded to the experimental group. The total number of PV+, PNN+ and PV+ neurons surrounded by PNNs (PV+PNN+) from the LA/BLA was estimated using a modified version of the fractionator method as described previously [59]. That is, all labeled cells covering the 100% of the LA/BLA area within each 50-μm-thick section of one from the six series of sections. The density of PV+, PNN+ and PV+PNN+ neurons per mm^3^ was calculated and statistics were performed using the number of animals as the sample number (n). The ratio of PV+ PNN+/PV+ neurons was also calculated and statistically analyzed as described below.

**Figure 2.**
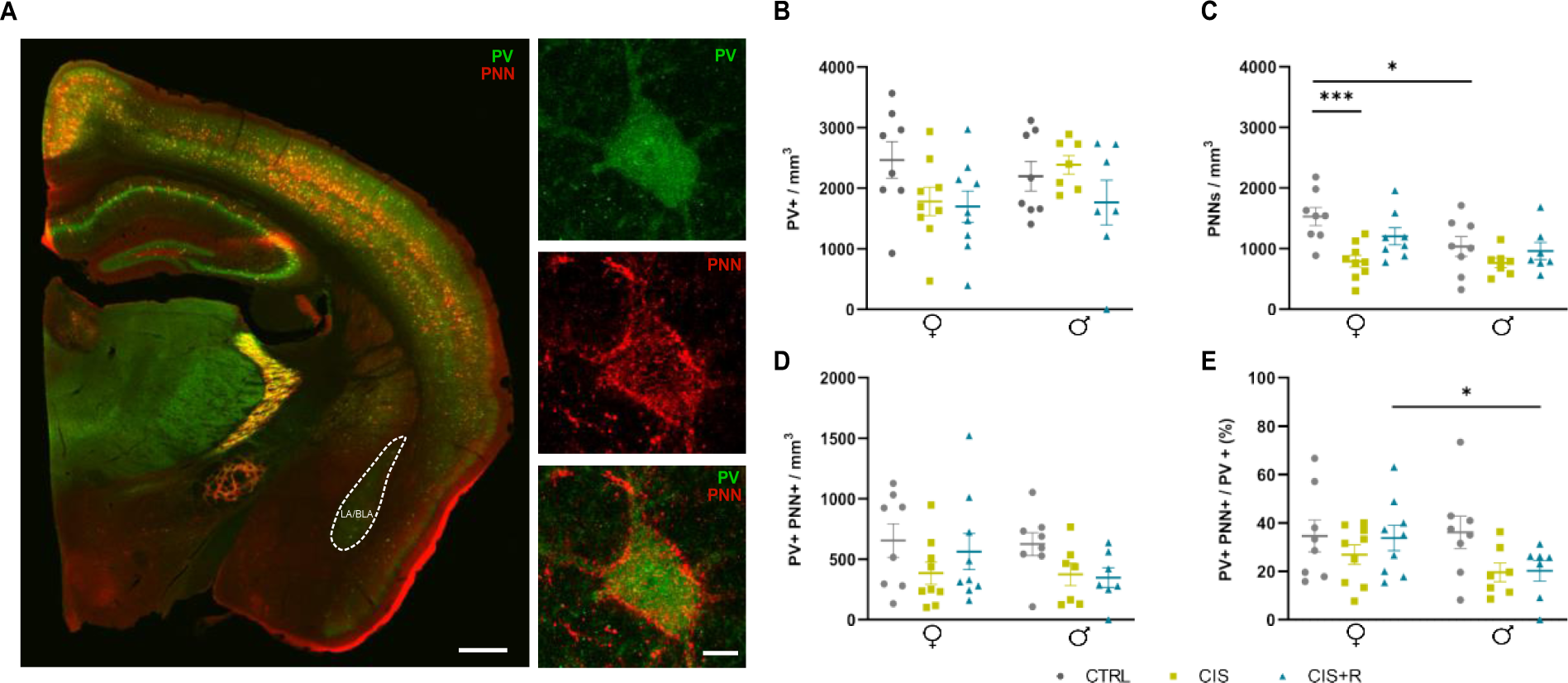
Analysis of parvalbumin (PV) expressing cells and perineuronal nets (PNNs) in the Lateral (LA) and Basolateral Amygdala (BLA). (A) Representative confocal images of brain sections immunostained for PV-expressing cells (green) and PNNs (red). For tile scan reconstruction (left panel), images were taken using a 10X objective, scale bar: 500 µm; and for PV+, PNN+ and PV+PNN+ pictures from LA/BLA (right panel), images were taken using 63X immersion objective with 4X digital, scale bar: 5µm. (B-E) Quantification of the density of PV+ and PNN+ neurons. Graphs represent the density of PV+ cells in LA/BLA complex as PV+/mm^3^ (B), density of PNNs as PNNs/mm^3^ (C), density of PV expressing cells surrounded by PNNs as PV+PNN+/mm^3^ (D) and the ratio of PV expressing cells surrounded by PNNs/PV expressing cells expressed as a percentage (E). Data in graphs are presented as mean ± SEM (n=7-9/group) and asterisks represent statistically significant differences among groups, p<0.05 (*), p<0.001 (***). Each dot represents each animal. PV, parvalbumin; PNN, perineuronal net; CTRL, Control group; CIS, Chronic isolation stress; CIS+R, Chronic isolation stress + resocialization.

#### Study of PNN integrity in the LA/BLA

For each animal, 6 PV+PNN+ neurons in the LA/BLA were randomly selected from two 50- µm-thick brain sections sampled with 250-µm distance (**Fig. 3A**). Z-stacks of confocal images covering the whole depth of the selected neuron were obtained with a 63x oil immersion objective and an additional 4x digital zoom with 0.20 μm step size (Leica SP8, Leica Microsystems, Wetzlar, Germany). To study the integrity of PNNs, we used a semi-automatic tracing method of the holes adapted from a previous work [39] with some modifications. In brief, 2D reconstructions from the z-stacks were obtained after applying a nonlinear smoothing filter to images for adaptive noise reduction with FIJI Software (ImageJ, Maryland, USA). Then, channels were separated and the background color threshold for each PNN was obtained. In the next step, the image was binarized (**Fig. 3A**) and holes were defined as the areas where the fluorescence intensity of the PNN (WFA) remains lower than the given threshold (**Fig. 3B-C**). Then, the program provided an automatic outline of the PNN (**Fig. 3A**). The mean hole size and the percentage of the area occupied by holes in relation to the somatic area was calculated for every neuron as indicators of PNN integrity (low integrity: bigger holes and increased % of somatic area occupied with holes; **Fig. 3A**). Mean ±SEM was calculated for every animal and statistics were performed using the number of animals as the *n*.

**Figure 3.**
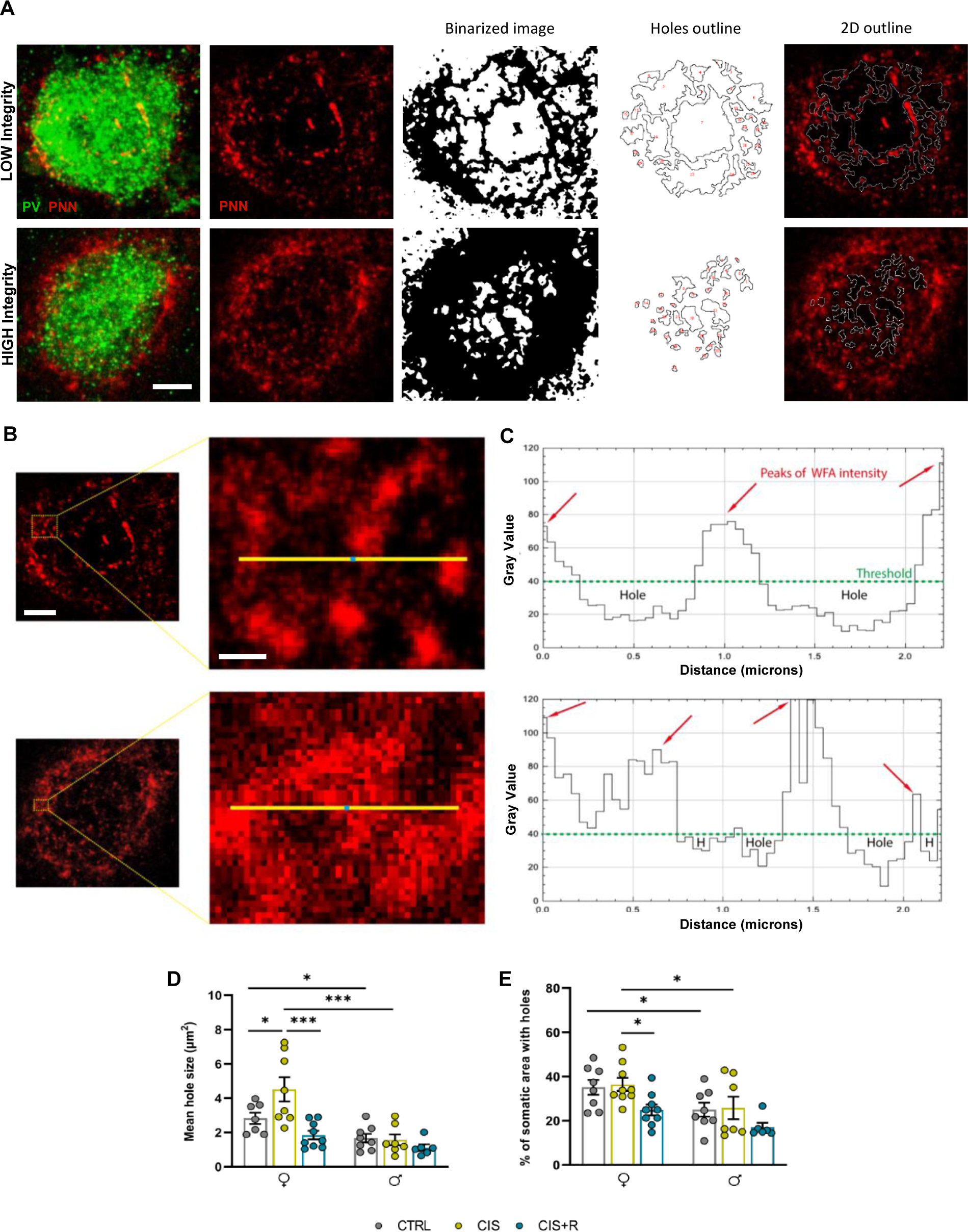
PNN integrity quantification. (A) Representative images of PNNs surrounding parvalbumin-expressing neurons in LOW integrity condition and HIGH integrity condition, and schematic representation of image processing for holes reconstruction and all PNN units traced by the program (binarized, holes outline and 2D outline). (B) Zoomed image of the yellow rectangular area from left panel, showing different holes size from the two different conditions. (C) The line profile of WFA intensity measured by Gray Value along the yellow line from panel B showing the peaks of WFA (red arrows) and the holes size measured by distance (microns). The dashed green line defines the threshold that was set to define holes. (D) Graph representing average size of PNNs holes (total size/number of holes) expressed in µm². (E) Percentage occupied by PNNs holes in the PV+ neuron soma. Scale bar: 5 µm (A and B left), 0,5 µm (B right). Each dot represents each animal. Data in graphs are presented as mean ± SEM (n=7-9/group) and asterisks represent statistically significant differences among groups, p<0.05 (*), p<0.001 (***). PV, parvalbumin; PNN, perineuronal net; CTRL, Control group; CIS, Chronic isolation stress; CIS+R, Chronic isolation stress + resocialization.

### Statistical Analysis

Statistical analysis was performed using GraphPad Prism 10.0 (GraphPad Software Inc., La Jolla, USA) and SPSS v28.0 (IBM Corp., NY, USA). After testing for normality using Shapiro-Wilk test, group differences were evaluated by means of two-way ANOVA with the number of animals as the *n*. When the effect of principal factors (sex or experimental condition) or interaction between them was statistically significant (p < 0.05), multiple pairwise comparisons between groups were performed (Bonferroni post hoc analyses). Data in the figures are expressed as mean ± SEM and p < 0.05 are indicated.

## Results

### Social isolation and successive resocialization modulate affective behavior in a sex-dependent manner

To assess whether social isolation contributes to the development of depression and anxiety-like symptoms in old mice and whether an equivalent regrouping period can rescue animals from these symptoms, we exposed our experimental females and males to a FST and OF procedure, respectively.

### Social isolation induces depressive-like symptoms in old females that are rescued by resocialization

For the FST, we evaluated the immobility time, episodes, and latency as indicatives of depressive-like behavior (**Fig. 1C**). Two-way ANOVA revealed a main effect of rearing in the time immobile (p=0.039) and in immobility episodes (p=0.002), but no effect was found in immobility latency (**Table 1**). Moreover, a significant “rearing x sex” interaction effect was found in all the 3 parameters measured in the FST (time immobile: p=0.007; immobility episodes: p=0.023; immobility latency: p=0.034; **Table 1**). Multipair wise comparison showed significantly decreased time immobile (p=0.008) and episodes (p=0.005) in CIS females when compared to CTRL group (**Fig. 1C**, **Table 2**). Remarkably, the alterations in these parameters reported in females were recovered after a resocialization period (CIS vs CIS+R: p=0.008 for immobility time and p=0.004 for immobile episodes; **Fig. 1C**, **Table 2**). No significant differences between CTRL males and CIS males were found in any of the parameters analyzed in the FST, suggesting that old males are more resilient than females to the effects of social isolation stress on depressive-like behavior (**Fig. 1C**, **Table 2**). In fact, CIS males showed decreased immobility time (p=0.003) and immobility episodes (p=0.010) than CIS females (**Fig. 1C**, **Table 2**). Interestingly, although CIS does not seem to affect depressive-like behavior in males, a resocialization period (CIS+R) decreased immobility episodes in CIS+R males when compared to their respective CTRL (p=0.026; **Fig. 1C**, **Table 2**)

**Table 1.** Summary of statistical results (I): Two-way ANOVA.

| Parameter |  | Main effects<br>(Two-way ANOVA) |  |  |
| --- | --- | --- | --- | --- |
| Behavior |  | Rearing | Sex | Interaction |
|  | Depressive |  |  |  |
| | Immobility time | $F_{2,42}=3.513; p=0.039 *$ | $F_{1,42}=1.916; p=0.174$ | $F_{2,42}=5.523; p=0.007 **$ |
| | Immobility episodes | $F_{2,42}=7.137; p=0.002 **$ | $F_{1,42}=3.347; p=0.074 \#$ | $F_{2,42}=4.147; p=0.023 *$ |
| | Immobility latency | $F_{2,42}=0.856; p=0.432$ | $F_{1,42}=0.147; p=0.703$ | $F_{2,42}=3.653; p=0.034 *$ |
|  | Locomotor Activity |  |  |  |
| | Distance travelled | $F_{2,42}=0.056; p=0.946$ | $F_{1,42}=0.206; p=0.652$ | $F_{2,42}=0.406; p=0.669$ |
|  | Anxiety |  |  |  |
| | Time Center / Time Periphery (10 min) | $F_{2,42}=0.086; p=0.917$ | $F_{1,42}=0.111; p=0.741$ | $F_{2,42}=1.302; p=0.283$ |
| | Time Center / Time Periphery (0-5 min) | $F_{2,40}=0.335; p=0.717$ | $F_{1,40}=0.623; p=0.435$ | $F_{2,40}=0.628; p=0.539$ |
| | Time Center / Time Periphery (5-10 min) | $F_{2,42}=0.086; p=0.917$ | $F_{1,42}=0.111; p=0.741$ | $F_{2,42}=1.302; p=0.283$ |
| Plasticity |  |  |  |  |
|  | Density (number cells/mm2) |  |  |  |
| | PV+ | $F_{2,42}=2.551; p=0.090 \#$ | $F_{1,42}=0.379; p=0.542$ | $F_{2,42}=1.351; p=0.270$ |
| | PNN+ | $F_{2,41}=7.445; p=0.002 **$ | $F_{1,41}=5.523; p=0.024 *$ | $F_{2,41}=1.586; p=0.217$ |
| | PV+ PNN+ | $F_{2,42}=2.698; p=0.079 \#$ | $F_{1,42}=0.810; p=0.373$ | $F_{2,42}=0.483; p=0.620$ |
| | % PV+ PNN+ / PV+ | $F_{2,42}=1.586; p=0.217$ | $F_{1,42}=2.524; p=0.120$ | $F_{2,42}=2.060; p=0.140$ |
|  | Integrity |  |  |  |
| | Mean hole size | $F_{2,39}=7.538; p=0.002 **$ | $F_{1,39}=24.315; p<0.001 ***$ | $F_{2,39}=4.338; p=0.020 *$ |
| | % somatic area with holes | $F_{2,41}=5.500; p=0.008 **$ | $F_{1,41}=12.381; p=0.001 **$ | $F_{2,41}=0.107; p=0.899$ |

Symbols and abbreviations: $p<0.05$ (\*), $p<0.01$ (\*\*), $p<0.001$ (\*\*\*), $0.10 \geq p \geq 0.05$ (#). PV, parvalbumin; PNN, perineuronal net.

**Table 2.** Summary of statistical results (II): multiple pairwise comparisons.

| Parameter |  | Multiple pairwise comparisons with Bonferroni's correction |  |  |  |  |  |  |  |  |
| --- | --- | --- | --- | --- | --- | --- | --- | --- | --- | --- |
|  |  | Sex comparisons |  |  | Female |  |  | Male |  |  |
| Behavior |  | CTRL | CIS | CIS+R | CTRL<br>*x*<br>CIS | CIS<br>*x*<br>CIS+R | CTRL<br>*x*<br>CIS+R | CTRL<br>*x*<br>CIS | CIS<br>*x*<br>CIS+R | CTRL<br>*x*<br>CIS+R |
|  | Depressive |  |  |  |  |  |  |  |  |  |
|  | Immobility time | *p*=0.119 | *p*=0.003<br>** | *p*=0.319 | *p*=0.008<br>** | *p*=0.008<br>** | *p*=1.000 | *p*=0.385 | *p*=1.000 | *p*=0.084<br># |
|  | Immobility episodes | *p*=0.229 | *p*=0.010<br>* | *p*=0.102 | *p*=0.005<br>** | *p*=0.004<br>** | *p*=1.000 | *p*=1.000 | *p*=0.145 | *p*=0.026<br>* |
|  | Immobility latency | *p*=0.073<br># | *p*=0.060<br># | *p*=0.584 | *p*=0.050 | *p*=0.275 | *p*=1.000 | *p*=0.578 | *p*=1.000 | *p*=0.387 |
|  | Locomotor Activity |  |  |  |  |  |  |  |  |  |
|  | Distance travelled | *p*=0.322 | *p*=0.924 | *p*=0.911 | *p*=1.000 | *p*=1.000 | *p*=1.000 | *p*=1.000 | *p*=1.000 | *p*=1.000 |
|  | Anxiety |  |  |  |  |  |  |  |  |  |
|  | Time Center /<br>Time Periphery<br>(10 min) | *p*=0.779 | *p*=0.327 | *p*=0.206 | *p*=1.000 | *p*=0.403 | *p*=1.000 | *p*=1.000 | *p*=1.000 | *p*=1.000 |
|  | Time Center /<br>Time Periphery<br>(0-5 min) | *p*=0.601 | *p*=0.227 | *p*=0.716 | *p*=1.000 | *p*=1.000 | *p*=1.000 | *p*=1.000 | *p*=0.685 | *p*=1.000 |
|  | Time Center /<br>Time Periphery<br>(5-10 min) | *p*=0.779 | *p*=0.327 | *p*=0.206 | *p*=1.000 | *p*=0.403 | *p*=1.000 | *p*=1.000 | *p*=1.000 | *p*=1.000 |
| Plasticity |  |  |  |  |  |  |  |  |  |  |
|  | Density (number<br>cells/mm2) |  |  |  |  |  |  |  |  |  |
|  | PV+ | *p*=0.481 | *p*=0.119 | *p*=0.861 | *p*=0.296 | *p*=1.000 | *p*=0.126 | *p*=1.000 | *p*=0.391 | *p*=0.816 |
|  | PNN+ | *p*=0.014<br>* | *p*=0.887 | *p*=0.438 | *P* <0.001<br>*** | *p*=0.083 # | *p*=0.254 | *p*=0.498 | *p*=0.979 | *p*=1.000 |
|  | PV+ PNN+ | *p*=0.860 | *p*=0.952 | *p*=0.195 | *p*=0.285 | *p*=0.750 | *p*=1.000 | *p*=0.430 | *p*=1.000 | *p*=0.318 |
|  | % PV+ PNN+ /<br>PV+ | *p*=0.613 | *p*=0.375 | *p*=0.023<br>* | *p*=1.000 | *p*=0.266 | *p*=0.752 | *p*=0.210 | *p*=1.000 | *p*=0.295 |
|  | Integrity |  |  |  |  |  |  |  |  |  |
|  | Mean hole size | *p*=0.021<br>* | *p*<0.001<br>*** | *p*=0.251 | *p*=0.014 * | *p*<0.001<br>*** | *p*=0.240 | *p*=1.000 | *p*=1.000 | *p*=1.000 |
|  | % somatic area<br>with holes | *p*=0.033<br>* | *p*=0.026<br>* | *p*=0.117 | *p*=1.000 | *p*=0.031 * | *p*=0.078 # | *p*=1.000 | *p*=0.289 | *p*=0.353 |

Symbols and abbreviations: $p < 0.05$ (\*), $p < 0.01$ (\*\*), $p < 0.001$ (\*\*\*), $0.10 \geq p \geq 0.05$ (#). PV, parvalbumin; PNN, perineuronal net; CTRL, Control group; CIS, Chronic isolation stress; CIS+R, Chronic isolation stress + resocialization.

### Anxiety-like behavior and locomotor activity are not affected by social isolation or resocialization in old mice, regardless of sex

In the OF test, (**Fig. 1D**) locomotor activity and anxiety-like behavior were evaluated, respectively, by means of the distance travelled by the animal (**Fig. 1E**) and the ratio between the time they spent in the center zone versus the time they spent in the periphery zone (TC/TP ratio; **Fig. 1F**). No statistically significant differences were observed in locomotor activity neither in males nor in females (**Fig. 1E, Tables 1&2**). In the same line, TC/TP ratio remain unchanged in all conditions and in both sexes (**Fig. 1F, Tables 1&2**). This ratio was also stable when considering the first 5 min of the test (**Fig. 1G, Tables 1&2**). As no effect of rearing was found in locomotor activity in the OF, the statistically significant impairments that old animals showed in the FST (see before) cannot be associated to locomotor impairments.

### Neuronal plasticity in the LA/BLA is affected by social isolation and subsequent resocialization but just in females

PNNs tightly control the plasticity [32–34] and stability of neuronal circuits in key brain regions for affective functions like the amygdala [27]. To investigate whether depressive-like symptoms caused by social isolation in old females are related to PNN impairments, we first quantified the density of PNNs in the LA/BLA in relation to PV+ neurons (**Fig. 2**) and afterwards analyzed PNN integrity (**Fig. 3**). Old males were also analyzed for comparative purposes.

### The density of PNNs in the LA/BLA of old females is reduced after social isolation and partially rescued by resocialization

Two-way ANOVA results (**Table 1**) showed no main effects of rearing, sex, or interaction in the density of PV+ neurons (**Fig. 2B**), the density of PV+PNN+ neurons (**Fig. 2D**) or the % of PV+ cells that co-express PNN (**Fig. 2E**). However, a main effect of rearing (p=0.002) and sex (p=0.024) in the density of PNNs was reported (**Table 1**). Specifically, CIS females showed lower density of PNNs (p<0.001) compared to CTRL and a trend towards a recovery was observed after resocialization (p=0.083). Interestingly, no statistically significant changes in the density of PNNs were observed in males, supporting our hypothesis that old males are more resilient to the effects of social isolation than old females (**Fig. 2C**, **Table 2**). In fact, CTRL males showed decreased PNN density when compared to CTRL females (p= 0.014; **Fig. 2C**, **Table 2**) and CIS+R males, decreased % of PV+ neurons that co-express PNN (p=0.023; **Fig. 2E**, **Table 2**) compared to CIS+R females.

### Social isolation disintegrates PNNs in the LA/BLA of old females, but resocialization rescues these impairments

The integrity of PNNs is regulated by the size of their holes [38, 39], in such a way that smaller holes stabilize existing synapses (high integrity and reduced plasticity) while larger holes leave “free” space for new synapses (low integrity and increased plasticity). In this study, we reconstructed, outlined, and automatically analyzed the holes of PNNs in the LA/BLA of our experimental animals (**Fig. 3 A-C**). Two-way ANOVA showed a significant effect of rearing (p=0.002), sex (p<0.001) and interaction (p=0.020) in the mean hole size (**Table 1**). Interestingly, females showed bigger holes after CIS (p=0.014) that decreased in size once animals were subjected to subsequent resocialization (p<0.001), suggesting that the plasticity of PV+ neurons in the LA/BLA is regulated by social environment, at least in old females (**Fig. 3D**, **Table 2**). In line with other parameters analyzed in our study, males did not show any alteration in the mean hole size of their PNNs, suggesting again an increased resilience of males to the effects of social isolation stress (**Fig. 3D**, **Table 2**). In fact, CTRL males showed decreased mean hole size (lower plasticity) when compared to CTRL females (p=0.021) and the same holds true for CIS males when compared to CIS females (p<0.001; **Fig. 3D**, **Table 2**). Interestingly, these results indicated differential basal levels of PNN integrity in males vs females and suggest that the lower plasticity of this region in old males might protect them from the affective effects of isolation stress.

The percentage of somatic area occupied by holes may also be indicative of altered integrity of PNNs, however other studies have demonstrated increased % of somatic area occupied by holes because of more numerous but smaller holes, thus indicative of higher but not lower integrity of PNNs [39]. In our study, two-way ANOVA showed significant effects of rearing (p=0.008) and sex (p=0.001) in the % of somatic area occupied by holes (**Table 1**). Nevertheless, multiple pairwise comparison only revealed significant changes when compared CIS vs CIS+R female groups (p=0.031; **Fig. 3E**, **Table 2**). In line with other results, CIS did not affect this parameter in male mice, reinforcing the hypothesis of increased resilience in males. Actually, CTRL males showed decreased % of somatic area occupied by holes when compared to CTRL females (p= 0.033) and the same holds true for CIS males when compared to CIS females (p=0.026; **Fig. 3E**, **Table 2**).

## Discussion

The prevalence of social isolation is dramatically high in the older human population [9, 10] and it has been associated with an increased risk of suffering from major depression [6, 11–14], especially in women [1–3]. However, while the psychological effects of social isolation have been extensively studied, the neurobiological basis of this association is complex and still poorly understood. Because exploring this vital topic in humans is difficult, rodent models devoid of cultural connotations represent consistent models for understanding how social isolation affects the brain. In these animal models, the effects of prolonged social isolation (>1 month), especially during early stages of life, have been exhaustively investigated [61]. However, very little is known about the neurobiological effects of shorter isolation periods (<1 month, referred here as “chronic isolation stress, CIS”), and even less at old age [62–64].

In the adult population, evidence from animal models and human studies indicates that plasticity, especially that of limbic brain regions like LA/BLA [19–23], is impaired in depression [15, 16] but, till date, scarce studies address this question in older population [17, 18]. Given the lockdown recommendations established almost worldwide due to the COVID-19 pandemics, understanding the effects of social isolation in old age has gained momentum and several clinical and preclinical studies have emerged since then [14, 64–67]. Nevertheless, no preclinical studies till date have investigated the relationship between neuronal plasticity in the LA/BLA and resilience to social isolation stress during aging. Moreover, little attention has been paid to whether old males and females respond differently to CIS and whether a successive resocialization period can rescue all putative impairments at the behavioral and cellular level [66].

Therefore, in the present study, we examined the impact of CIS and resocialization in aged male and female mice, focusing our attention on affective symptoms (anxiety and depression) and the plasticity of PV+ neurons in the LA/BLA. We found that, in old females, CIS induced depressive-like symptoms and impaired LA/BLA plasticity by decreasing the density and the integrity of PNN. Importantly, resocialization effectively rescued all these impairments. By contrast, old males were not affected by CIS. Indeed, under social conditions, old males showed higher PNN integrity than females in LA/BLA, suggesting lower basal levels of plasticity for males (**Fig. 4**). In the following paragraphs we will discuss these results in comparison with previous preclinical and clinical studies.

**Figure 4.**
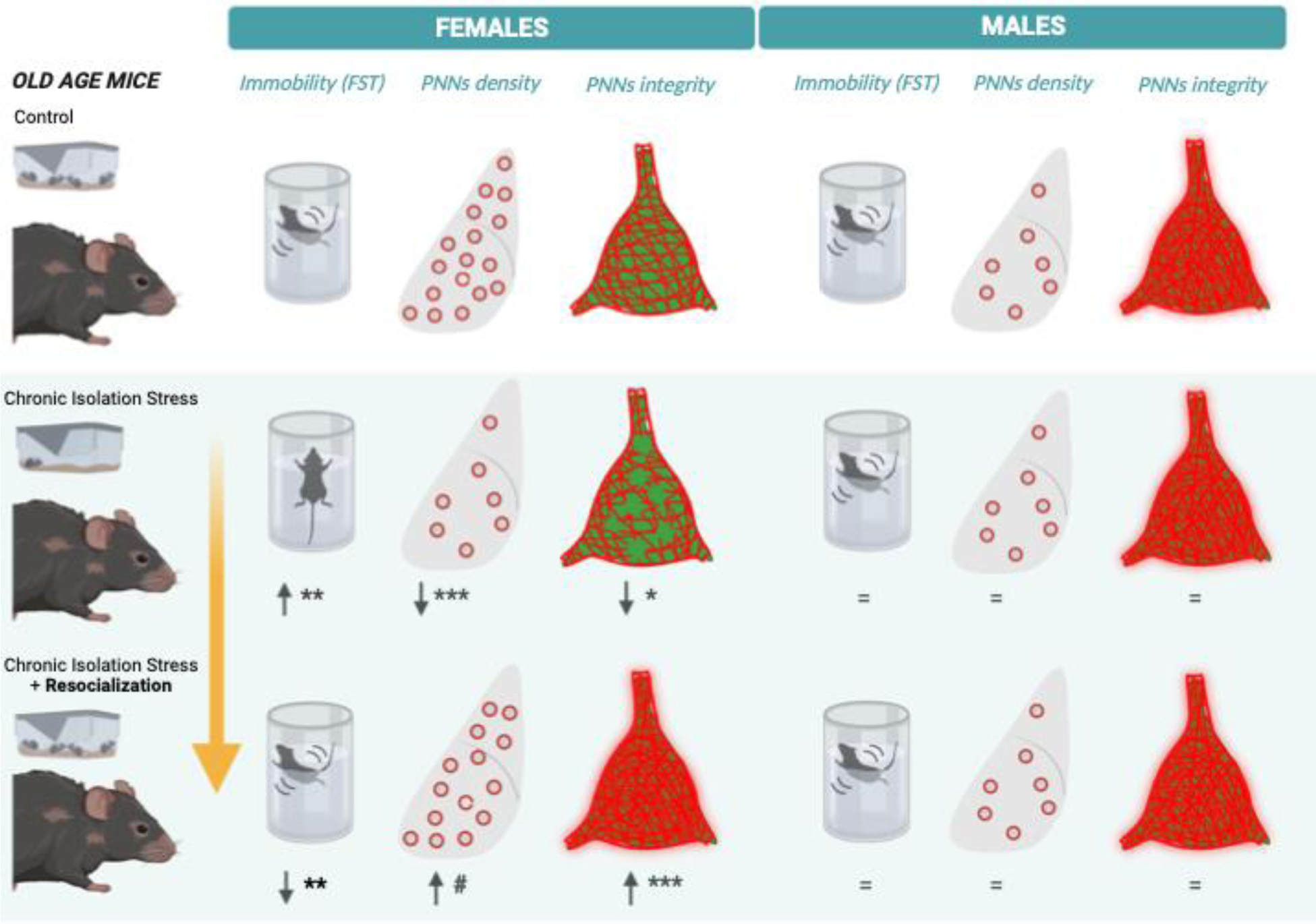
Graphical summary of results. In the present study, we investigated the impact of chronic isolation stress (CIS) and a subsequent resocialization period in aged male and female mice (∼ 21 months-old), focusing our attention on affective symptoms and the plasticity of parvalbumin-expressing (PV+) neurons in the lateral/basolateral amygdala (LA/BLA). We found that CIS impaired affective behaviour and LA/BLA plasticity only in females. Specifically, CIS induced depressive-like symptoms and decreased the integrity of perineuronal nets (PNN). Resocialization effectively rescued all these impairments. Note that old males were not affected by CIS but in CTRL conditions showed higher PNN integrity (less plasticity) than females. Symbols: p<0.05 (*), p<0.01 (**), p<0.001 (***), 0.10 ≥ p ≥ 0.05 (#). PV, parvalbumin; PNN, perineuronal net; CTRL, Control group; CIS, Chronic isolation stress; CIS+R, Chronic isolation stress + resocialization. Drawings created with BioRender.com.

Our experimental old females showed decreased immobility time and episodes in the FST after social isolation. These parameters are commonly used indicators of depressive-like behavior in rodents [55]. By contrast, old male mice demonstrated no effects of CIS when tested in the FST, suggesting that old males are more resilient than females to the behavioral consequences of social isolation stress (**Fig. 4**). Although many studies have reported that long-term isolation rearing induces depressive-like behavior in young and adult rodents, independently of sex [68–71], only few studies have addressed this question in old female [64] and male [72] mice. Our results are in agreement with the study of Sullens *et al.* that reported a mild effect of CIS on depressive-like behavior in 18 month-old female mice [64] and with results of Panossian *et al.* [72] showing no effects in 24 month-old male mice. Interestingly, as we commented before, depression has also been frequently described after a social isolation period but especially in aged women [11–14, 73]. Taken together, these findings suggest that social isolation might always be deleterious for females in terms of depression-like symptoms, whereas a lack of response to this stress can be expected for males at late-life stage. Remarkably, our study is the first preclinical study describing a recovery of depressive-like symptoms after a resocialization period in old females (**Fig. 4**), although this therapy has been effectively validated in isolated rodents [66, 74] and humans [75, 76] for improving other behavioral deficits associated to social isolation. In this line, it is woth noting that, although in our study CIS does not seem to affect depressive-like behavior in males, a resocialization period (CIS+R) decreased immobility episodes in CIS+R males when compared to their respective CTRL, suggesting a beneficial effect of resocialization also for males.

By contrast, our OF results demonstrated no effect of social isolation or resocialization on anxiety and locomotor activity in old mice, independently of sex. To date, limited studies have evaluated these parameters in aged rodents, with only two studies, to our knowledge, evaluating anxiety and locomotion after at least two weeks of isolation in females [64] and males [72], and no studies evaluating resocialization. Our results are in accordance with those described in 24 month-old male mice [72]. By contrast, Sullens *et al.* [64] reported hyperlocomotion and a mild anxiolytic effect in 18 month-old females induced after 4 weeks of social isolation, in consonance with other studies conducted in adolescent male mice [59, 77, 78]. The age of onset and/or the duration of the isolation protocol might justify the inconsistency of results between our study and Sullens’ [64]. Actually, OF results from other studies in adult mice show a great discrepancy in the effects of social isolation on anxiety and locomotion with some studies describing no effects, as we reported here, and some others showing conflicting results ([68] for review).

Several studies have demonstrated that PNNs downregulate neuronal plasticity [32–34] in critical brain areas for affective functioning, such as de amygdala, by creating a physical barrier that prevents the creation of new synapses while enhancing the stability of those that already exist [27, 35–38]. Indeed, a growing body of evidence suggests that PNNs in the amygdala are sensitive structures to the impact of stress, which breaks this “protective barrier” of PV+ neurons and enhance circuit instability [5]. Although several experimental studies indicate that the effects of stress on PNNs vary with age and brain region [5, 49], no preclinical research has yet examined this issue in the LA/BLA of elderly mice. Our study is the first describing a sex-dependent decrease of PNN density in this region after social isolation stress and a subsequent mild recovery induced by resocialization in old mice (**Fig. 4**). Specifically, and in accordance with behavioral data discussed above, females but not males were the ones affected, supporting again the hypothesis that old females are less resilient to the effects of social isolation than old males. As PNN associated neurons in the amygdala of rodents express estrogen receptors [79], a role for female hormones mediating this decreased resiliency to the effects of CIS is feasible. Indeed, this hypothesis is in agreement with studies suggesting that female estrogen fluctuations may be a trigger for depression ([80] for review).

PNNs are disintegrated by ECM remodeling enzymes, specially the MMP-9 that is mainly synthetized by neurons and glial cells [34, 40]. A recent report, suggests that the integrity of PNNs is regulated by the size of their holes; smaller holes stabilize existing synapses (reduced plasticity) while larger holes promote the formation of new synapses (increased plasticity) [39]. PNNs in our experimental females showed bigger holes after CIS and, importantly, its hole size was recovered after resocialization, suggesting that PNN integrity in the LA/BLA is regulated by social environment, at least in old females (**Fig. 4**). One possibility is that female hormones like estrogens regulate the integrity level of PNN in the amygdala by regulating MMP-9 levels, as it has been demonstrated that 17β-estradiol treatment *in vitro* increases MMP-9 levels released in the medium by neuroblastoma cultivated cells [81]. In line with other results in our study, males from our study did not show any alteration in PNN integrity, suggesting again an increased resilience of males to the effects of social isolation stress. Interestingly, we report here that PNN integrity in the LA/BLA of old males is higher than that of old females, suggesting that the lower plasticity of this region in old males might protect them from the affective effects of isolation stress. The percentage of somatic area occupied by holes can also be indicative of altered integrity of PNNs, however other studies have demonstrated increased % of somatic area occupied by holes because of more numerous but smaller holes, thus indicative of higher but not lower integrity of PNNs [39]. Indeed, in our study, CTRL males showed decreased % of somatic area occupied by holes when compared to CTRL females.

In summary, our study demonstrates that old females are more vulnerable to CIS than old males but can recover from depressive symptoms by resocialization. Moreover, the present results point to the integrity of PNN in the LA/BLA as a key regulator of depressive-like symptoms induced by social isolation. Therefore, our study can help on the understanding of the neurobiological link between social isolation and depression in aged population and might be important in the development of more effective treatments for depression, especially in old women.

## Acknowledgments

Funding: This research was financially supported by the European Commission (MSCA- SE101086247), Spanish Ministry of Science, Innovation and Universities (RTI2018-095698-B-I00), Generalitat Valenciana (AICO/2021/246) and Universitat Jaume I (UJI-A2020- 20). Author contributions: E.C-G designed the study. A.M-O and E.C-G run the experiment, behaviorally tested the animals, and perfused them. J.H-C cut the brains and analyzed the vaginal smears. J.H-C and A.M-O performed immunohistochemical labeling and confocal imaging. J.H- C analyzed the behavioral tests and confocal images. J-H-C and C.G-M performed the statistical analysis. J.H-C, A.M-O and C-G.M designed the figures and illustrations and wrote the initial draft of the manuscript. E.C-G wrote the final version of the manuscript. F.O-B reviewed and improved the manuscript. E.C-G and F.O-B edited the manuscript and figures. All authors read and approved the final manuscript

## Competing interests

Authors declare that they do not have any financial interests that could be perceived as a conflict of interest.

## Notes

### Competing Interest Statement

The authors have declared no competing interest.

## References

1. Tetsuka S (2021). Depression and Dementia in Older Adults: A Neuropsychological Review. Aging Dis, 12:1920–1934.

2. Luppa M, Sikorski C, Luck T, Ehreke L, Konnopka A, Wiese B, et al. (2012). Age- and gender-specific prevalence of depression in latest-life--systematic review and meta-analysis. J Affect Disord, 136:212–221.

3. Girgus JS, Yang K, Ferri C V. (2017). The Gender Difference in Depression: Are Elderly Women at Greater Risk for Depression Than Elderly Men? Geriatrics. doi: 10.3390/GERIATRICS2040035.

4. Mañas-Ojeda A, Ros-Bernal F, Olucha-Bordonau FE, Castillo-Gómez E (2020). Becoming Stressed: Does the Age Matter? Reviewing the Neurobiological and Socio-Affective Effects of Stress throughout the Lifespan. Int J Mol Sci 2020, Vol 21, Page 5819, 21:5819.

5. Perez-Rando M, Carceller H, Castillo-Gomez E, Bueno-Fernandez C, García-Mompó C, Gilabert-Juan J, et al. (2022). Impact of stress on inhibitory neuronal circuits, our tribute to Bruce McEwen. Neurobiol Stress, 19:100460.

6. Hussenoeder FS, Conrad I, Pabst A, Luppa M, Stein J, Engel C, et al. (2022). Different Areas of Chronic Stress and Their Associations with Depression. Int J Environ Res Public Health. doi: 10.3390/ijerph19148773.

7. Zapater-Fajarí M, Crespo-Sanmiguel I, Pulopulos MM, Hidalgo V, Salvador A (2021). Resilience and Psychobiological Response to Stress in Older People: The Mediating Role of Coping Strategies. Front Aging Neurosci, 13:67.

8. Dye C, Boerma T, Evans D, Harries A, Lienhardt C, McManus J, et al. The world health report 2013: research for universal health coverage. WHO Press: Geneva, Switzerland; 2013.

9. Yang K, Victor C (2011). Age and loneliness in 25 European nations. Ageing Soc, 31:1368–1388.

10. National Academies of Sciences E (2020). Social Isolation and Loneliness in Older Adults: Opportunities for the Health Care System. Soc Isol Loneliness Older Adults. doi: 10.17226/25663.

11. Noguchi T, Saito M, Aida J, Cable N, Tsuji T, Koyama S, et al. (2021). Association between social isolation and depression onset among older adults: a cross-national longitudinal study in England and Japan. BMJ Open. doi: 10.1136/BMJOPEN-2020-045834.

12. Leigh-Hunt N, Bagguley D, Bash K, Turner V, Turnbull S, Valtorta N, et al. (2017). An overview of systematic reviews on the public health consequences of social isolation and loneliness. Public Health, 152:157–171.

13. Santini ZI, Koyanagi A, Tyrovolas S, Mason C, Haro JM (2015). The association between social relationships and depression: A systematic review. J Affect Disord, 175:53–65.

14. Cacioppo JT, Hughes ME, Waite LJ, Hawkley LC, Thisted RA (2006). Loneliness as a specific risk factor for depressive symptoms: Cross-sectional and longitudinal analyses. Psychol Aging, 21:140.

15. Albert PR (2019). Adult neuroplasticity: A new “cure” for major depression? J Psychiatry Neurosci, 44:147–150.

16. Rădulescu I, Drăgoi AM,, Trifu SC, Cristea MB (2021). Neuroplasticity and depression: Rewiring the brain’s networks through pharmacological therapy (Review). Exp Ther Med. doi: 10.3892/ETM.2021.10565.

17. Bhandari A, Lissemore JI, Rajji TK, Mulsant BH, Cash RFH, Noda Y, et al. (2018). Assessment of Neuroplasticity in Late-Life Depression with Transcranial Magnetic Stimulation. J Psychiatr Res, 105:63.

18. Sibille E (2013). Molecular aging of the brain, neuroplasticity, and vulnerability to depression and other brain-related disorders. Dialogues Clin Neurosci, 15:53–65.

19. Adolphs R (2010). What does the amygdala contribute to social cognition? Ann N Y Acad Sci, 1191:42–61.

20. Adolphs R, Tranel D, Damasio H, Damasio A (1994). Impaired recognition of emotion in facial expressions following bilateral damage to the human amygdala. Nature, 372:669–672.

21. Hintiryan H, Bowman I, Johnson DL, Korobkova L, Zhu M, Khanjani N, et al. (2021). Connectivity characterization of the mouse basolateral amygdalar complex. Nat Commun 2021 121, 12:1-25.

22. Zhang X, Ge TT, Yin G, Cui R, Zhao G, Yang W (2018). Stress-induced functional alterations in amygdala: Implications for neuropsychiatric diseases. Front Neurosci, 12:367.

23. Liu W, Ge T, Leng Y, Pan Z, Fan J, Yang W, et al. (2017). The Role of Neural Plasticity in Depression: From Hippocampus to Prefrontal Cortex. Neural Plast, 2017:1–11.

24. Scheimann JR, Mahbod P, Morano R, Frantz L, Packard B, Campbell K, et al. (2018). Deletion of glucocorticoid receptors in forebrain GABAergic neurons alters acute stress responding and passive avoidance behavior in female mice. Front Behav Neurosci, 12:325.

25. Johnson LR, Farb C, Morrison JH, McEwen BS, Ledoux JE (2005). Localization of glucocorticoid receptors at postsynaptic membranes in the lateral amygdala. Neuroscience, 136:289–299.

26. Woodruff AR, Sah P (2007). Networks of Parvalbumin-Positive Interneurons in the Basolateral Amygdala. J Neurosci, 27:553.

27. Lupori L, Totaro V, Cornuti S, Ciampi L, Carrara F, Grilli E, et al. (2023). A comprehensive atlas of perineuronal net distribution and colocalization with parvalbumin in the adult mouse brain. Cell Rep, 42:112788.

28. Celio MR, Spreafico R, De Biasi S, Vitellaro-Zuccarello L (1998). Perineuronal nets: past and present. Trends Neurosci, 21:510–515.

29. Brückner G, Szeöke S, Pavlica S, Grosche J, Kacza J (2006). Axon initial segment ensheathed by extracellular matrix in perineuronal nets. Neuroscience, 138:365–375.

30. Celio MR, Blumcke I (1994). Perineuronal nets--a specialized form of extracellular matrix in the adult nervous system. Brain Res Brain Res Rev, 19:128–145.

31. Favuzzi E, Marques-Smith A, Deogracias R, Winterflood CM, Sánchez-Aguilera A, Mantoan L, et al. (2017). Activity-Dependent Gating of Parvalbumin Interneuron Function by the Perineuronal Net Protein Brevican. Neuron, 95:639–655.e10.

32. van ’t Spijker HM, Kwok JCF (2017). A sweet talk: The molecular systems of perineuronal nets in controlling neuronal communication. Front Integr Neurosci, 11:33.

33. Sorg BA, Berretta S, Blacktop JM, Fawcett JW, Kitagawa H, Kwok JCF, et al. (2016). Casting a Wide Net: Role of Perineuronal Nets in Neural Plasticity. J Neurosci, 36:11459–11468.

34. Fawcett JW, Oohashi T, Pizzorusso T (2019). The roles of perineuronal nets and the perinodal extracellular matrix in neuronal function. Nat Rev Neurosci 2019 208, 20:451- 465.

35. Hensch TK (2004). Critical period regulation. Annu Rev Neurosci, 27:549–79.

36. Hensch TK, Bilimoria PM (2012). Re-opening Windows: Manipulating Critical Periods for Brain Development. Cerebrum, 2012:11.

37. Wang D, Fawcett J (2012). The perineuronal net and the control of CNS plasticity. Cell Tissue Res, 349:147–60.

38. Carceller H, Guirado R, Ripolles-Campos E, Teruel-Marti V, Nacher J (2020). Perineuronal nets regulate the inhibitory perisomatic input onto parvalbumin interneurons and gamma activity in the prefrontal cortex. J Neurosci, 40:5008–5018.

39. Kaushik R, Lipachev N, Matuszko G, Kochneva A, Dvoeglazova A, Becker A, et al. (2021). Fine structure analysis of perineuronal nets in the ketamine model of schizophrenia. Eur J Neurosci, 53:3988–4004.

40. Reinhard SM, Razak K, Ethell IM (2015). A delicate balance: Role of MMP-9 in brain development and pathophysiology of neurodevelopmental disorders. Front Cell Neurosci, 9:280.

41. Murase S, Lantz CL, Quinlan EM (2017). Light reintroduction after dark exposure reactivates plasticity in adults via perisynaptic activation of MMP-9. Elife. doi: 10.7554/ELIFE.27345.

42. Foscarin S, Raha-Chowdhury R, Fawcett JW, Kwok JCF (2017). Brain ageing changes proteoglycan sulfation, rendering perineuronal nets more inhibitory. Aging (Albany NY), 9:1607–1622.

43. Yang S, Gigout S, Molinaro A, Naito-Matsui Y, Hilton S, Foscarin S, et al. (2021). Chondroitin 6-sulphate is required for neuroplasticity and memory in ageing. Mol Psychiatry 2021 2610, 26:5658-5668.

44. Hernandes-Alejandro M, Montaño S, Harrington CR, Wischik CM, Salas-Casas A, Cortes-Reynosa P, et al. (2020). Analysis of the Relationship Between Metalloprotease-9 and Tau Protein in Alzheimer’s Disease. J Alzheimer’s Dis, 76:553–569.

45. Bitanihirwe BKY, Woo TUW (2020). A conceptualized model linking matrix metalloproteinase-9 to schizophrenia pathogenesis. Schizophr Res, 218:28–35.

46. Li H, Sheng Z, Khan S, Zhang R, Liu Y, Zhang Y, et al. (2022). Matrix Metalloproteinase-9 as an Important Contributor to the Pathophysiology of Depression. Front Neurol, 13:861843.

47. Wen TH, Binder DK, Ethell IM, Razak KA (2018). The Perineuronal ’Safety’ Net? Perineuronal Net Abnormalities in Neurological Disorders. Front Mol Neurosci, 11:270.

48. Carceller H, Gramuntell Y, Klimczak P, Nacher J (2022). Perineuronal Nets: Subtle Structures with Large Implications: https://doi.org/101177/10738584221106346, 10738584221106346.

49. Laham BJ, Gould E (2022). How Stress Influences the Dynamic Plasticity of the Brain’s Extracellular Matrix. Front Cell Neurosci, 15:574.

50. Nelson JF, Felicio LS, Osterburg HH, Finch CE (1981). Altered profiles of estradiol and progesterone associated with prolonged estrous cycles and persistent vaginal cornification in aging C57BL/6J mice. Biol Reprod, 24:784–794.

51. Felicio LS, Nelson JF, Finch CE (1984). Longitudinal studies of estrous cyclicity in aging C57BL/6J mice: II. Cessation of cyclicity and the duration of persistent vaginal cornification. Biol Reprod, 31:446–453.

52. Caligioni CS (2009). Assessing Reproductive Status/Stages in Mice. Curr Protoc Neurosci. doi: 10.1002/0471142301.nsa04is48.

53. Ajayi AF, Akhigbe RE (2020). Staging of the estrous cycle and induction of estrus in experimental rodents: an update. Fertil Res Pract 2020 61, 6:1-15.

54. Cora MC, Kooistra L, Travlos G (2015). Vaginal Cytology of the Laboratory Rat and Mouse: Review and Criteria for the Staging of the Estrous Cycle Using Stained Vaginal Smears. Toxicol Pathol, 43:776–793.

55. Can A, Dao DT, Arad M, Terrillion CE, Piantadosi SC, Gould TD (2011). The Mouse Forced Swim Test. J Vis Exp. doi: 10.3791/3638.

56. Fitzgerald PJ, Yen JY, Watson BO (2019). Stress-sensitive antidepressant-like effects of ketamine in the mouse forced swim test. PLoS One, 14:e0215554.

57. Bueno-Fernandez C, Perez-Rando M, Alcaide J, Coviello S, Sandi C, Castillo-Gómez E, et al. (2021). Long term effects of peripubertal stress on excitatory and inhibitory circuits in the prefrontal cortex of male and female mice. Neurobiol Stress, 14:100322.

58. Perez-Rando M, Castillo-Gomez E, Bueno-Fernandez C, Nacher J (2018). The TrkB agonist 7,8-dihydroxyflavone changes the structural dynamics of neocortical pyramidal neurons and improves object recognition in mice. Brain Struct Funct. doi: 10.1007/s00429-018-1637-x.

59. Castillo-Gómez E, Pérez-Rando M, Bellés M, Gilabert-Juan J, Llorens JV, Carceller H, et al. (2017). Early social isolation stress and perinatal nmda receptor antagonist treatment induce changes in the structure and neurochemistry of inhibitory neurons of the adult amygdala and prefrontal cortex. eNeuro. doi: 10.1523/ENEURO.0034-17.2017.

60. Härtig W, Brauer K, Brückner G (1992). Wisteria floribunda agglutinin-labelled nets surround parvalbumin-containing neurons. Neuroreport, 3:869–872.

61. Mumtaz F, Khan MI, Zubair M, Dehpour AR (2018). Neurobiology and consequences of social isolation stress in animal model-A comprehensive review. Biomed Pharmacother, 105:1205–1222.

62. Shoji H, Mizoguchi K (2011). Aging-related changes in the effects of social isolation on social behavior in rats. Physiol Behav, 102:58–62.

63. Wang L, Cao M, Pu T, Huang H, Marshall C, Xiao M (2018). Enriched Physical Environment Attenuates Spatial and Social Memory Impairments of Aged Socially Isolated Mice. Int J Neuropsychopharmacol, 21:1114–1127.

64. Sullens DG, Gilley K, Jensen K, Vichaya E, Dolan SL, Sekeres MJ (2021). Social isolation induces hyperactivity and exploration in aged female mice. PLoS One, 16:e0245355.

65. Manca R, De Marco M, Venneri A (2020). The Impact of COVID-19 Infection and Enforced Prolonged Social Isolation on Neuropsychiatric Symptoms in Older Adults With and Without Dementia: A Review. Front psychiatry. doi: 10.3389/FPSYT.2020.585540.

66. Lavenda-Grosberg D, Lalzar M, Leser N, Yaseen A, Malik A, Maroun M, et al. (2021). Acute social isolation and regrouping cause short- and long-term molecular changes in the rat medial amygdala. Mol Psychiatry 2021 272, 27:886-895.

67. Brooks SK, Webster RK, Smith LE, Woodland L, Wessely S, Greenberg N, et al. (2020). The psychological impact of quarantine and how to reduce it: rapid review of the evidence. Lancet, 395:912–920.

68. Grigoryan GA, Pavlova I V., Zaichenko MI (2022). Effects of Social Isolation on the Development of Anxiety and Depression-Like Behavior in Model Experiments in Animals. Neurosci Behav Physiol 2022 525, 52:722-738.

69. Berry A, Bellisario V, Capoccia S, Tirassa P, Calza A, Alleva E, et al. (2012). Social deprivation stress is a triggering factor for the emergence of anxiety- and depression-like behaviours and leads to reduced brain BDNF levels in C57BL/6J mice. Psychoneuroendocrinology, 37:762–772.

70. Takatsu-Coleman AL, Patti CL, Zanin KA, Zager A, Carvalho RC, Borçoi AR, et al. (2013). Short-term social isolation induces depressive-like behaviour and reinstates the retrieval of an aversive task: Mood-congruent memory in male mice? J Psychiatry Neurosci, 38:259–268.

71. Liu N, Wang Y, An AY, Banker C, Qian YH, O’Donnell JM (2020). Single housing- induced effects on cognitive impairment and depression-like behavior in male and female mice involve neuroplasticity-related signaling. Eur J Neurosci, 52:2694–2704.

72. Panossian A, Cave MW, Patel BA, Brooks EL, Flint MS, Yeoman MS (2020). Effects of age and social isolation on murine hippocampal biochemistry and behavior. Mech Ageing Dev, 191:111337.

73. Courtin E, Knapp M (2017). Social isolation, loneliness and health in old age: a scoping review. Health Soc Care Community, 25:799–812.

74. An D, Chen W, Yu DQ, Wang SW, Yu WZ, Xu H, et al. (2017). Effects of social isolation, re-socialization and age on cognitive and aggressive behaviors of Kunming mice and BALB/c mice. Anim Sci J, 88:798–806.

75. Hoang P, King JA, Moore S, Moore K, Reich K, Sidhu H, et al. (2022). Interventions Associated With Reduced Loneliness and Social Isolation in Older Adults. JAMA Netw Open, 5:e2236676.

76. Piolatto M, Bianchi F, Rota M, Marengoni A, Akbaritabar A, Squazzoni F (2022). The effect of social relationships on cognitive decline in older adults: an updated systematic review and meta-analysis of longitudinal cohort studies. BMC Public Health, 22:1–14.

77. Fone KCF, Porkess MV (2008). Behavioural and neurochemical effects of post-weaning social isolation in rodents-relevance to developmental neuropsychiatric disorders. Neurosci Biobehav Rev, 32:1087–102.

78. Dimonte S, Sikora V, Bove M, Morgese MG, Tucci P, Schiavone S, et al. (2023). Social isolation from early life induces anxiety-like behaviors in adult rats: Relation to neuroendocrine and neurochemical dysfunctions. Biomed Pharmacother, 158:114181.

79. Ciccarelli A, Weijers D, Kwan W, Warner C, Bourne J, Gross CT (2021). Sexually dimorphic perineuronal nets in the rodent and primate reproductive circuit. J Comp Neurol, 529:3274–3291.

80. Albert PR (2015). Why is depression more prevalent in women? J Psychiatry Neurosci, 40:219–221.

81. Merlo S, Sortino MA (2012). Estrogen activates matrix metalloproteinases-2 and -9 to increase beta amyloid degradation. Mol Cell Neurosci, 49:423–429.

